# Alpha cell dysfunction in early type 1 diabetes

**DOI:** 10.1101/2021.10.15.464545

**Authors:** Nicolai M Doliba, Andrea V Rozo, Jeffrey Roman, Wei Qin, Daniel Traum, Long Gao, Jinping Liu, Elisabetta Manduchi, Chengyang Liu, Maria L Golson, Golnaz Vahedi, Ali Naji, Franz M Matschinsky, Mark A. Atkinson, Alvin C Powers, Marcela Brissova, Klaus H Kaestner, Doris A Stoffers, for the HPAP Consortium

## Abstract

Multiple islet autoantibodies (AAb) predict type 1 diabetes (T1D) and hyperglycemia within 10 years. By contrast, T1D develops in just ∼15% of single AAb+ (generally against glutamic acid decarboxylase, GADA+) individuals; hence the single GADA+ state may represent an early stage of T1D amenable to interventions. Here, we functionally, histologically, and molecularly phenotype human islets from non-diabetic, GADA+ and T1D donors. Similar to the few remaining beta cells in T1D islets, GADA+ donor islets demonstrated a preserved insulin secretory response. By contrast, alpha cell glucagon secretion was dysregulated in both T1D and GADA+ islets with impaired glucose suppression of glucagon secretion. Single cell RNA sequencing (scRNASeq) of GADA+ alpha cells revealed distinct abnormalities in glycolysis and oxidative phosphorylation pathways and a marked downregulation of *PKIB*, providing a molecular basis for the loss of glucose suppression and the increased effect of IBMX observed in GADA+ donor islets. The striking observation of a distinct early defect in alpha cell function that precedes beta cell loss in T1D suggests that not only overt disease, but also the progression to T1D itself, is bihormonal in nature.

## Introduction

Longitudinal studies have shown that individuals at high genetic risk for or with a family history of T1D who later develop diabetes progress through several distinct stages prior to the onset of clinical symptoms (Insel et al., 2015). The presence of islet AAb is currently the best biomarker for the future onset of hyperglycemia in T1D. The presence of two or more AAb confers a 70% risk of T1D developing within 10 years and nearly 100% over the lifetime of the individual. The factors involved in the rate of progression are poorly understood, although a younger age at seroconversion, a higher number of positive AAb, and higher levels of IAA and IA-2A AAb have been associated with a more rapid rate of progression to T1D (So et al., 2021; Steck and Rewers, 2011; Ziegler et al., 2013). In contrast, T1D develops in just 15% of single AAb+ individuals within 10 years of follow-up, with anti-glutamic acid decarboxylase AAb (GADA) by far the most common presenting AAb (Steck and Rewers, 2011; Ziegler et al., 2013). T1D is typically characterized by the progressive loss of insulin-secreting beta cells (Insel et al., 2015). However, while largely underappreciated, glucagon-secreting alpha cells are also affected in T1D and contribute to the pathophysiology of diabetes (Christensen et al., 2011; Emami et al., 2017; Kulina and Rayfield, 2016; Lee et al., 2016). Indeed, in T1D islets, alpha cell function is compromised while the few remaining beta cells are functionally nearly normal (Brissova et al., 2018).

Glucagon is the main secretory product of pancreatic alpha cells. The function of this peptide hormone is to increase glucose production and thus provide sustained glucose supply to the brain and other vital organs during fasting conditions. Thus, targeting of the pancreatic alpha cell and its main secretory product glucagon has potential as treatment for diabetes (Christensen et al., 2011). Glucagon secretion from alpha cells is impacted by cAMP (Yu et al., 2019) and Ca^2+^ (Vieira et al., 2007), as well as intra-islet insulin (Xu et al., 2006), GABA (Kittler and Moss, 2003; Xu et al., 2006), cAMP-activated ion channels (Huang et al., 2017), and gap junctions (Dufer, 2018). A direct role for glucose sensing via glucokinase in alpha cells has also been identified as a key factor in glucose inhibition of glucagon secretion (Bahl et al., 2021; Basco et al., 2018; Moede et al., 2020). In addition, glucagon released from alpha cells acts in a paracrine fashion on beta cells, with this crosstalk important for high efficiency glucose stimulation of insulin secretion (Sharma et al., 2018; Svendsen et al., 2018; Zhu et al., 2019b). This said, our understanding of the progression of beta and alpha-cell dysfunction during the development of T1D is incomplete, in part due to the difficulty in obtaining T1D pancreata and islets for functional analysis.

The NIDDK-funded Human Pancreas Analysis Program (HPAP; https://hpap.pmacs.upenn.edu) provides comprehensive physiological and molecular profiling of the pancreas during T1D pathogenesis and publicly disseminates simultaneous analyses of human islet physiology, metabolism, molecular profiling, and immunobiology from well characterized deceased donors (Kaestner et al., 2019). Utilizing donor tissues and cells from HPAP, we examined both insulin and glucagon secretion from non-diabetic, AAb+, and T1D islets. We make the striking observation of a distinct early defect in alpha cell function that precedes beta cell loss during the progression of T1D, a finding that suggests that not only overt disease, but also the progression to T1D itself, is bihormonal in nature.

## Results and Discussion

### Glucose Sensitivity is Preserved in the Remaining Beta Cells but Lost in Alpha Cells of Islets from T1D Donors

We performed insulin and glucagon secretion studies by perifusion of islets isolated from 18 normoglycemic organ donors with no autoantibodies to islet antigens (‘control’), 9 normoglycemic organ donors positive for AAb against GAD (‘GADA+’) and 6 organ donors with T1D (Supplemental Table S1). Both basal and maximal rates of insulin secretion were dramatically reduced in T1D islets (1/60 that of normal islets) (Figure 1A). Remarkably, however, amino acids with low glucose, high glucose and IBMX all stimulated insulin secretion by T1D islets in a similar pattern to normal islets (Figure 1A). Total insulin release under high glucose stimulation (Figure 1B) and IBMX potentiation (Figure 1C) were 0.95% and 1.47%, respectively, that of control islets. Insulin content was markedly reduced in T1D islets as expected (Supplemental Figure S1A). Thus, while the baseline and maximal insulin secretion rates are dramatically reduced in T1D islets, the response of the few remaining beta cells to glucose and to cAMP elevation was preserved.

**Figure 1.**
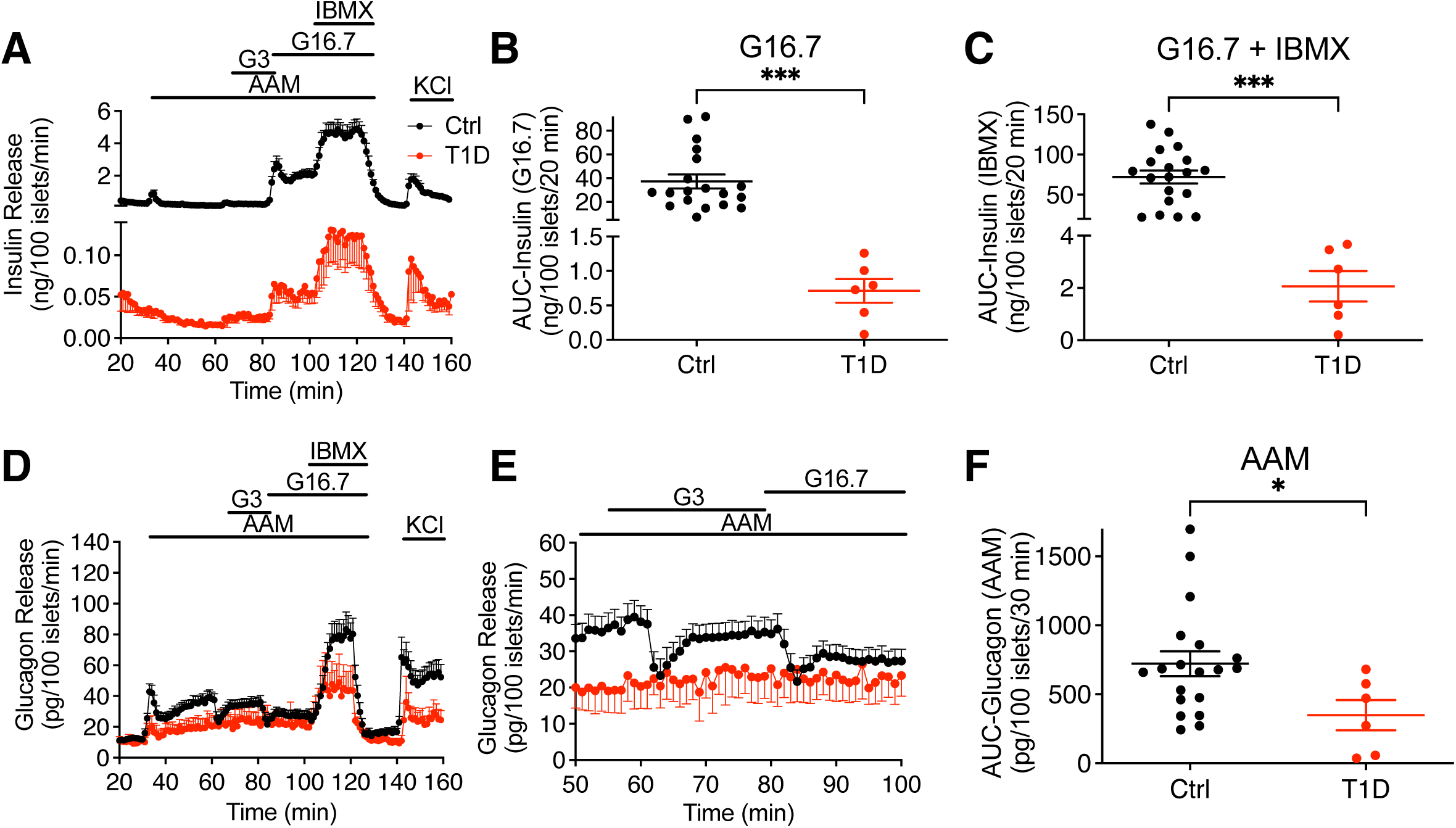
Insulin and glucagon secretion in islets from normal and T1D donors. (A) The dynamics of insulin secretion in response to different stimuli. (B) Total insulin secretion during 16 mM glucose-stimulation; (C) Total insulin secretion during IBMX-potentiation. The basal and maximal insulin secretion in T1D islets was 1/60 that of normal islets. Notably, T1D islets had similar insulin response pattern at low, high glucose, and IBMX compared to control islets. (D) Glucagon secretion profiles (E) Magnified view of selected section (53-100 min) of the experiment presented in Panel D to highlight the difference in glucose suppression of glucagon secretion between normal and T1D islets. (F) Total glucagon secretion during AAM-stimulation.

Both first and second phases of amino acid-stimulated glucagon secretion were significantly reduced in T1D islets (Figure 1D and 1F). Additionally, T1D islets lacked suppression of glucagon secretion by low and high glucose (Figure 1E). However, T1D islets showed no difference in IBMX-potentiated glucagon secretion (Figure S1E), while depolarization by KCl caused less glucagon secretion in T1D islets. Previously, preservation of stimulated insulin secretion and dysfunction of alpha cells in T1D donors was noted (Brissova et al., 2018). Our data indicate that, in addition to lower glucagon secretion, glucose does not suppress glucagon secretion. Thus, in striking contrast to the remarkably preserved insulin secretory response in remaining beta cells, the ability of glucose to inhibit glucagon secretion is lost in T1D islets.

### Glucose Suppression of Glucagon Secretion is Impaired in Islets from GADA+ Donors

Next, we utilized the nine HPAP islet preparations from single GADA+, normoglycemic individuals to investigate potential early alterations in islet function. Islets from the same preparations were analyzed simultaneously at Penn and at Vanderbilt using two complementary perifusion protocols to maximize the information that could be obtained and to cross-validate our findings. The protocol developed by Brissova and colleagues was previously employed for functional phenotyping of T1D islets (Brissova et al., 2018) and has been adopted by the Human Islet Phenotyping Program for functional analysis of human islet preparations made available for research by the Integrated Islet Distribution Program (https://iidp.coh.org/). The second protocol employed at Penn (see Methods) was specifically designed to be sensitive to changes in the alpha cell response to amino acids and low glucose. At both laboratories, stimulated insulin secretion profiles were similar between GADA+ and control cases (Figures 2A, S2A, and S2E-H).

**Figure 2.**
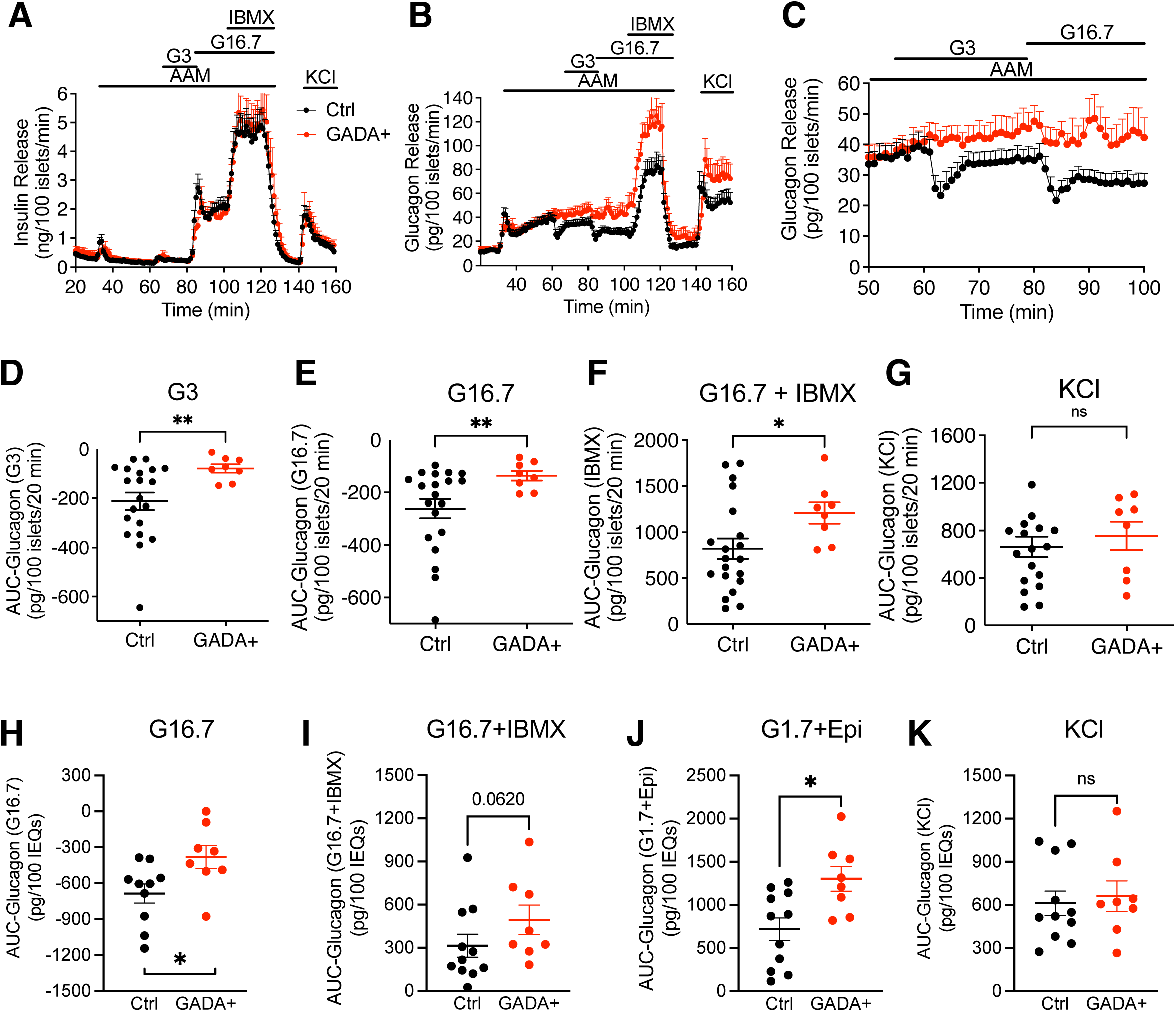
Insulin and glucagon secretion in islets from normal and GADA+ donors. (A) The dynamics of insulin secretion during different interventions. (B) The dynamics of glucagon secretion. (C) Magnified view of selected section (53-100 min) of the data from panel B highlights the difference in glucose suppression of glucagon secretion between normal and GADA+ islets. (D-G) Total glucagon secretion during 3 mM glucose (D), 16.7 mM glucose (E), G16.7 + IBMX (F), and KCl (G) calculated as area under curve (AUC). (H-K) Islets from the same preparations were assessed by perifusion assay at Vanderbilt University (see Figures S2A and S2B). AUC analysis of glucagon responses to high glucose (H) c-AMP-mediated secretion in response to IBMX (I) and epinephrine (J) and an unaltered KCl response (K) *p<0.05.

In striking contrast to insulin secretion, glucagon secretion was significantly altered in GADA+ donors. Amino acids induced biphasic glucagon secretion to the same extent in control and GADA+ islets, with the second phase of glucagon secretion monotonically increasing during amino acid stimulation in both groups of islets (Figure 2B). Low and high glucose caused sustained suppression of glucagon secretion in controls as expected. Surprisingly, however, in GADA+ donor islets, low glucose effected only a slight delay in the monotonically increasing second phase of glucagon secretion, while high glucose caused little to no suppression of glucagon secretion (Figure 2C). Area under the curve quantification showed that the suppression of glucagon release during low and high glucose was significantly lower in GADA+ donors (Figures 2D, 2E, 2H). In addition, potentiation of glucagon secretion by IBMX, a phosphodiesterase inhibitor that maximally increases intracellular cAMP concentrations, was ∼50% greater in islets from GADA+ donors compared to controls (Figure 2F). There was no difference in the readily releasable pool of glucagon granules (Figure 2G). Analysis of the same islet preparations with the Vanderbilt protocol confirmed decreased suppression of glucagon secretion by glucose, increased response to low glucose plus epinephrine and a trend toward increased response to IBMX, with no change in the KCl effect ((Figures S2B and 2H-2K). Insulin and glucagon content were also measured in all islet preparations; both insulin and glucagon content were similar between control and GADA+ donor islets. Together these data indicate that although GADA+ donors are normoglycemic as reflected by a normal A1c and have normal insulin secretion, glucagon secretion is notably altered.

### Islet Composition is not altered in GADA+ Donors

A difference in islet composition could explain the altered glucagon secretion in islets from GADA+ organ donors. To evaluate this possibility, we took advantage of the highly quantitative endocrine cell composition data available through HPAP. Firstly, we determined the proportions of endocrine cells in donor pancreata by flow CyTOF of single islet cell suspensions stained simultaneously with a panel of 36 antibodies. In contrast to our previous observation of an increased alpha cell fraction in T1D donors compared with controls (Wang et al., 2019), flow CYTOF analysis of 9 non-diabetic GADA+ and 12 controls revealed no statistically significant change in the alpha cell fraction in GADA+ islets (Figures 3A-C, S1C and S1D). Secondly, we analyzed data from imaging mass cytometry (IMC) performed on the same donor; these images were segmented into thousands of individual cells, which were then identified by their protein marker expression. Again, as shown in Figures 3D and S3A, there was no statistically significant difference in endocrine cell populations between the two groups. Figures 3E and 3F show representative multi-channel overlays of pancreatic islets from control and GADA+ organ donors that illustrate this point. To assess whether immunological infiltrates in or near islets are already present at the single GADA+ stage, we quantified immune cell infiltrates from our IMC images and performed detailed image analysis of the distance relationships between pancreatic islets and CD4+ and CD8+ T cells, proliferating (activated) and non-proliferating macrophages, B cells, and Treg cells but found no significant differences (Figure S4).

**Figure 3.**
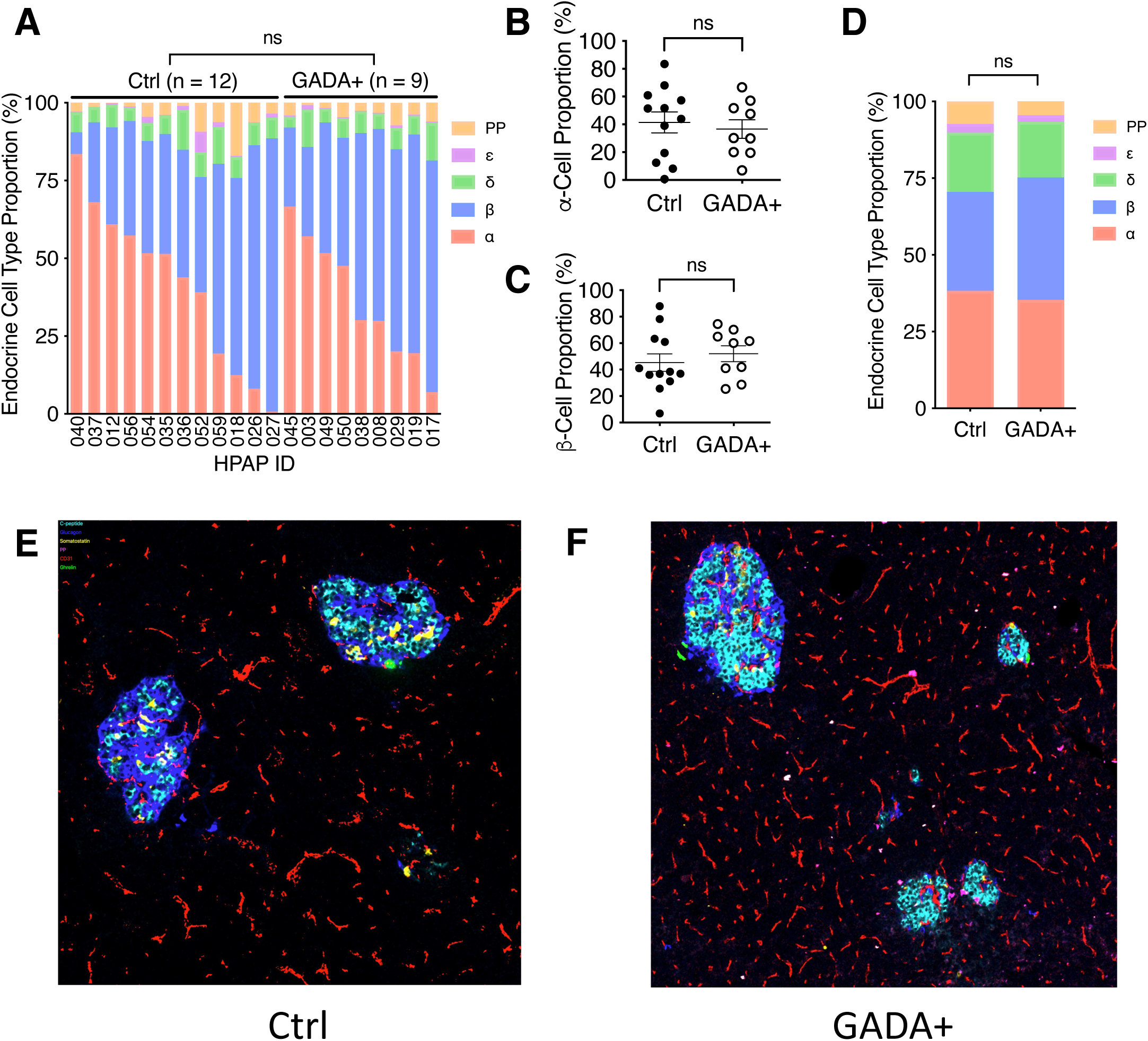
Islet composition of control and GADA+ donors. (A-C) Endocrine cell type proportions determined by Flow CyTOF of single cell suspensions of islets. (A) Cell type fraction plotted for the indicated HPAP cases. (B,C) Average alpha (B) and beta (C) cell percentage as determined by Flow CyTOF. (D) Endocrine cell type proportions determined by IMC. (E,F) Representative examples of IMC images of control (E) and GADA+ (F) pancreas with 7 channels shown, green, insulin; red, glucagon; yellow, somatostatin; orange, PECAM; white, Pancreatic polypeptide; magenta, ghrelin; purple, CD68 (macrophages).

### Transcriptome Alterations in Alpha Cells from GADA+ Donors

Given that islet endocrine cell composition, islet architecture, and immune cell infiltration were not significantly altered in GADA+ donors, we next focused on alpha cell transcriptome changes to understand if they explain the observed alpha cell dysfunction. We analyzed scRNAseq data from control and GADA+ organ donors produced by HPAP, and post cell type clustering selected only cells in the major alpha cell cluster (data not shown). Next, we collapsed the transcriptome of all single alpha cells for each donor into ‘pseudobulk’ alpha cell transcriptomes and performed differential gene expression analysis. Overall, the alpha cell transcriptomes of the two groups were quite similar; however, specific genes and pathways were differentially expressed as shown in Figure 4. Panel A shows a heatmap of 52 differentially expressed genes identified by DESeq2. Most notable among the differentially expressed genes is *PKIB*, encoding the cAMP-dependent protein kinase inhibitor beta, which acts as competitive inhibitor of protein kinase A (Figure 4 and Table S2). *PKIB* transcripts are decreased on average 4.9 fold in alpha cells from GADA+ organ donors, suggesting a possible activation of the cAMP pathway consistent with the IBMX and epinephrine effect on glucagon secretion reported above.

**Figure 4.**
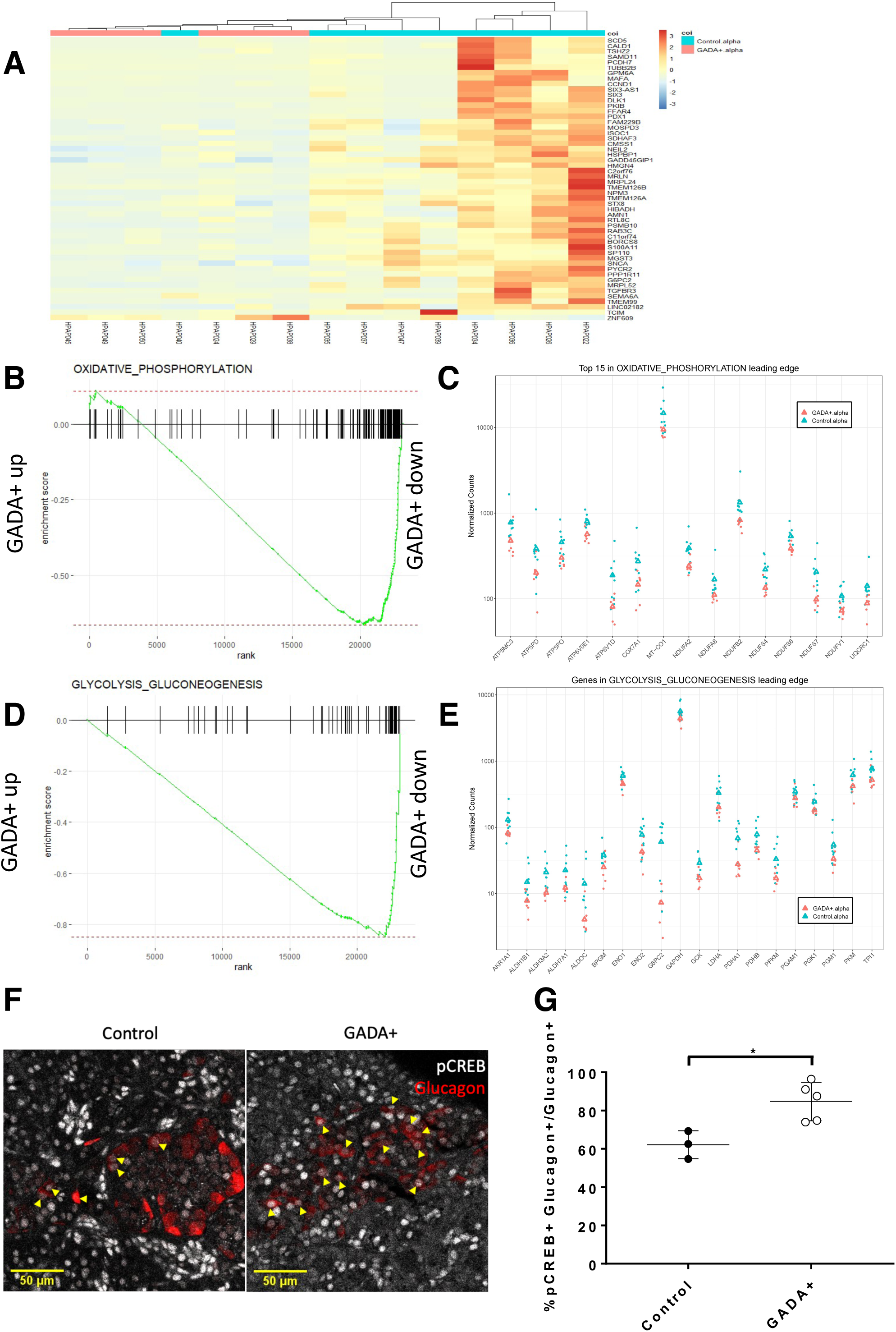
Single cell transcriptome analysis and immunofluorescence staining of control and GADA+ alpha cells. (A) Heatmap of hierarchical clustering of differentially expressed genes obtained by comparing pseudobulk alpha cell gene expression of 9 control and 6 GADA+ organ donors. This heatmap was generated with the R package pheatmap (coi = cells of interest). (B) Genes annotated to the ‘oxidative phosphorylation’ pathway are highly enriched among the genes downregulated in alpha cells from GADA+ organ donors. (C) Expression of the top 15 genes in the leading edge of the oxidative phosphorylation pathway. Triangles indicate means. (D) Genes annotated to the ‘glycolysis_gluconeogenesis’ pathway are highly enriched among the genes downregulated in alpha cells from GADA+ organ donors. (E) Expression of the genes in the leading edge of the ‘glycolysis_gluconeogenesis’ pathway. Triangles indicate means. (F) Immunofluorescence staining of pancreatic sections with antisera against pCREB and glucagon. Arrows indicate pCREB+ glucagon+ cells (G) Nuclear phosphoCREB positive alpha cells are increased in frequency in GADA+ islets. (n=3 for control and 5 for GADA+).

Next, we employed gene set enrichment analysis (GSEA), a method that detects statistically significant changes in transcript levels for preselected groups of functionally linked genes. One of the most significantly affected pathways was that of oxidative phosphorylation, which was enriched among genes with lower transcript levels in alpha cells from GADA+ organ donors (Figure 4B; adjusted p-value 0.008). The leading edge of this pathway (i.e., the core group of genes that accounts for the gene set’s enrichment signal) contains 75 genes, the top 15 of which are shown in Figure 4C. Among these genes are three that encode subunits of the mitochondrial ATP synthase as well as seven that encode subunits of the NADH:ubiquinone oxidoreductase complex (Complex 1), which may indicate a reduction in mitochondrial oxidative ATP synthesis. The glycolysis_gluconeogenesis pathway was also enriched among genes with lower transcript levels in alpha cells from GADA+ organ donors (Figure 4D; adjusted p-value 0.0002). The genes in the leading edge of this pathway are shown in Figure 4E. Remarkably, the entire set of glycolysis genes, from *GCK* to *PDH* is downregulated in alpha cells from GADA+ donors. It was recently demonstrated that the rate of glycolytic flux via glucokinase and thus ATP production determines the setpoint for inhibition of glucagon secretion by glucose (Bahl et al., 2021; Basco et al., 2018; Moede et al., 2020). Therefore, if GADA+ alpha cells produce ATP from glucose less efficiently because of lower glycolytic flux and oxidative phosphorylation, then a given concentration of glucose would less efficiently suppress glucagon secretion, providing a possible molecular explanation for the physiological observations shown in Figure 2.

The increased potentiation of glucagon secretion by IBMX and epinephrine described above (Figures 2B, S2B, 2F, 2I and 2J) suggested an upregulation of cAMP signaling pathway in alpha cells from GADA+ donors. As introduced above, we identified *PKIB*, a competitive inhibitor of protein kinase A, as the most differentially expressed in the alpha cells of GADA+ islets, providing a possible molecular explanation for this effect. To provide further evidence for altered cAMP signaling in alpha cells of GADA+ individuals, we performed immunofluorescence staining of human islet sections using anti-phospho CREB (P-CREB) antisera. Elevated cAMP stimulates the activity of protein kinase A which in turn phosphorylates the transcription factor CREB. The percent of alpha cells that expressed nuclear P-CREB was significantly elevated in GADA+ islets (Figures 4F and 4G). The defects in glucose suppression of glucagon secretion and elevation of cAMP signaling in GADA+ islets may be interrelated, since the glucose inhibition of glucagon secretion can be overcome by maintaining cAMP at high levels (Yu et al., 2019).

Historically, the development of diabetes has been characterized by defects in beta cells. Through our phenotyping of non-diabetic, AAb+ islets, we have discovered that alpha cell dysfunction precedes beta cell loss. This study highlights the need to consider T1D as a disease in which both beta and alpha cells are impacted. Future studies will be needed to expand on the mechanisms suggested above through the lenses of alpha cell physiology, immune signaling, and paracrine effects. In addition, given that only a subset of single AAb+ cases progress to T1D, further investigation on a larger number of AAb+ cases is needed to define the possible distinctions between likely progressors and non-progressors (Anand et al., 2021). With a comprehensive understanding of the role of the alpha cell in diabetes, a more targeted intervention may be developed at the critical single AAb+ stage.

## Acknowledgments

We thank Mike Feldman for providing sections of HPAP donor pancreata. We thank the donors and their families for their priceless contribution of human tissue. We acknowledge the PENN DRC (P30 DK019525) that houses the Penn Metabolic and Physiological Phenotyping Core and the Cellular and Molecular Phenotyping Core of the HPAP program which generated many of the datasets employed in this investigation. This work is supported by 3UC4-112217-01S1 and U01-DK123594-02, UC4-DK-112217, UC4-DK-112232, and U01-DK-123716.

## Author Contributions

ND, FM, MB, KK and DS were responsible for Conceptualization. ND, AR, WQ, DT, and JL were responsible for Investigation. ND, AR, DT, JL, EM, and JR were responsible for Visualization. ND, JR, DS and KK were responsible for Writing-Original Draft Preparation. Writing – Review & Editing Preparation was performed by MA, ACP, MB, KK and DS. Formal Analysis was conducted by JR, LG, EM and GV. Project Administration was carried out by MG. Critical Resources were provided by CL and AN. MB was responsible for Validation. Funding Acquisition was carried out by AN, MA, ACP, KK, MB and DS. MB, KK and DS were responsible for Supervision.

---

**Supplemental Figure S1**. (A) Insulin and (B) glucagon content of islets recovered after perifusion. Proportion of β (C) and α cells (D) in human islet preparations from controls (n=19), GADA+ (n=10) and T1D individuals (n=8). (E) Total glucagon secretion during IBMX-potentiation in islets from control and T1D donors. Data was analyzed by one way ANOVA with Tukey’s multiple comparisons test; *** p<0.001, **** p<0.0001.

**Supplemental Figure S2. Secretory profiles of islets from GADA+ organ donors and controls assessed at Vanderbilt University**. Dynamic insulin (A) and glucagon secretory response to various secretagogues measured by perifusion of control (Ctrl) and GADA+ islets; G 5.6 – 5.6 mM glucose; G 16.7 – 16.7 mM glucose; G 16.7 + IBMX 100 – 16.7 mM glucose with 100 μM isobutylmethylxanthine (IBMX); G1.7 + Epi 1 – 1.7 mM glucose and 1μM epinephrine; KCl 20 – 20 mM potassium chloride (KCl) was normalized to islet volume expressed by islet equivalents (IEQs); 1 IEQ corresponds to an islet with a diameter of 150 μm. (C) Islet insulin content. (D) Islet glucagon content. (E-H) Integrated area under the curve (AUC) analyses to insulin secretagogues highlighted in gray in panel A. * p<0.05; ** p<0.01. Error bars indicate SEM. Panels C-H were analyzed by two-tailed t-test.

**Supplemental Figure S3. Quantification of cell type proportions for all donors, determined by IMC**. Panels represent proportions of each endocrine cell type as a proportion of (A) total cell number and (B) the endocrine cells.

**Supplemental Figure S4. Immune cell infiltration as determined by imaging mass cytometry (IMC)**. For various immune cell types, cell distributions by distance-from-islet were determined. Violin plots of mean distance by cell type and donor are also presented. P-values for comparison between GADA+ donors and controls are shown in the lower-left of each panel.

**Supplemental Table S1. Donor Information**. *ICRH91 (UNOS ADBD275); ICRH99 (UNOS ADGB379); ICRH100 (UNOS ADID386)

**Supplemental Table S2. Differentially expressed genes between GADA+ and control alpha cells**. Gene expression data obtained by scRNAseq was compared between alpha cells from GADA+ and control donors. Genes with adjusted p-value (padj)<0.05 from the DESeq2 analyses are listed here. Fold change (FC) compares GADA+ donors to controls.

## STAR Methods RESOURCE AVAILABILITY

### Lead contact

Further information and requests for resources and reagents should be directed to and will be fulfilled by the lead contact, Doris Stoffers.

### Materials Availability

This study did not generate new unique reagents.

### Data and code availability

This paper analyzes existing, publicly available data generated by the authors. Raw image data for imaging mass cytometry, detailed clinical information, solution CyTOF as well as transcriptome and perifusion data are available from PancDB (https://hpap.pmacs.upenn.edu). Any additional information required to reanalyze the data reported in this paper is available from the lead contact upon request

## EXPERIMENTAL MODEL AND SUBJECT DETAILS

### Human Islets

GADA+ autoantibody-positive organ donors were screened by the nPOD/HPAP team as previously described (Wasserfall and Atkinson, 2006). Human islets were isolated from donor pancreata using standard multiorgan recovery and modified Ricordi techniques (Ricordi et al., 2016) at the accredited Human Islet Resource Center at the University of Pennsylvania (hereafter termed Penn). Following 2-3 days of culture in supplemented CMRL-1066 medium (Deng et al., 2004; Deng et al., 2003), we characterized the physiology and metabolic state of 39 human islet preparations isolated from 23 control (20 from HPAP, 3 from Penn’s Human Islet Resource Center; age 27 ± 10 years, BMI 27 ± 7 kg/m^2^, HbA1c 5.3 ± 0.6 [mean ± SD]), 10 GADA+ (age 24 ± 5 years, BMI 27 ± 4 kg/m^2^, HbA1c 5.4 ± 0.2) and 6 T1D-donors (age 20 ± 8 years, BMI 19 ± 5 kg/m^2^, HbA1c 10.1 ± 0.6). To age-match controls and those with GADA+ and T1D, only donors with age less than 40 years were included in this study. An aliquot of most islet preparations was also shipped to Vanderbilt for parallel analyses. The Penn Institutional Review Board considers this research exempt because islets were received from deceased, de-identified organ donors. All pancreata were acquired from deceased donors after obtaining consent from their families through UNOS (United Network for Organ Sharing)(Health Resources and Services Administration, 2020)

## METHOD DETAILS

### Perifusion of human islets

HPAP employs two complementary and validated islet perifusion protocols to assess insulin and glucagon secretion (at Penn and at Vanderbilt). At Penn, islets were pre-perifused with substrate-free medium. Next, a physiological amino acid mixture (AAM; mixture of 19 amino acids (Li et al., 2004); total concentration 4 mM) was added to stimulate glucagon secretion. Then, low and high glucose (3 and 16.7 mM) were added to stimulate insulin secretion and to inhibit glucagon secretion. During the high glucose step, IBMX (0.1 mM) was added to maximally increase intracellular cAMP levels and stimulate secretion of both hormones. Finally, a brief washout period with substrate-free medium removed all stimulants, and then 30 mM KCl was added to depolarize the islet cells and quantify the readily releasable pool of secretory granules. Insulin and glucagon in perifusion aliquots and insulin content were measured as previously described (Doliba et al., 2017). Glucagon content was measured by ELISA (Crystal Chem).

At Vanderbilt, islet function was assessed by perifusion as previously described (Brissova et al., 2018) and this methodology was adopted by the NIDDK-funded Human Islet Phenotyping Program (HIPP) of the Integrated Islet Distribution Program (https://www.protocols.io/view/analysis-of-islet-function-in-dynamic-cell-perifus-bt9knr4w). Insulin and glucagon concentration in perifusates and islet extracts was measured by radioimmunoassay (insulin, RI-13K; glucagon, GL-32K; Millipore).

### scRNASeq analysis

The Single Cell 3’ Reagent Kit v2 or v3 was used for generating scRNA-seq data. 3,000 cells were targeted for recovery per donor. All libraries were validated for quality and size distribution using a BioAnalyzer 2100 (Agilent) and quantified using Kapa (Illumina). For samples prepared using ‘The Single Cell 3’ Reagent Kit v2’, the following chemistry was performed on an Illumina HiSeq4000: Read 1: 26 cycles, i7 Index: 8 cycles, i5 index: 0 cycles, and Read 2: 98 cycles. For samples prepared using ‘The Single Cell 3’ Reagent Kit v3’, the following chemistry was performed on an Illumina HiSeq 4000: Read 1: 28 cycles, i7 Index: 8 cycles, i5 index: 0 cycles, and Read 2: 91 cycles. Cell Ranger (10x Genomics; v3.0.1) was used for bcl2fastq conversion, aligning (using the hg38 reference genome), filtering, counting, cell calling, and aggregating (--normalize=none) (fig. S1, a and b). scRNASeq data were preprocessed as described (Fasolino et al., 2021). We downloaded the resulting Seurat (Butler et al., 2018; Stuart et al., 2019) object from the cellxgene resource at https://cellxgene.cziscience.com/collections/51544e44-293b-4c2b-8c26-560678423380. Exclusion of donor HPAP-019 (whose scRNASeq library was from sorted beta cells) and donors younger than 5 or older than 40, resulted in 9 healthy control donors and 6 GADA+ donors (Suppl. Table 1). Working with R version 4.1.1, we generated pseudobulk raw counts for each gene by aggregating (via sum) counts of the alpha cells at the donor level. Raw counts were then input into DESeq2 (Love et al., 2014) for pre-processing and differential expression analysis with a p-value threshold of 0.05. Gene Set Enrichment Analysis (GSEA) (Mootha et al., 2003; Subramanian et al., 2005) was performed using the fgsea Bioconductor package (Sergushichev, 2016) with gene ranking based on the shrunken log2 Fold Changes derived using the apeglm Bioconductor package (Zhu et al., 2019a). We used the KEGG canonical pathway collection (c2.cp.kegg.v7.4.symbols.gmt) from MSigDB (Subramanian et al., 2005) augmented with the cAMP-Signaling pathway directly extracted from KEGG (Kanehisa and Goto, 2000), for a total of 187 gene sets.

### Flow CyTOF

The flow CyTOF experiments were performed as described (Wang et al., 2016). Briefly, after dissociated cells were barcoded following the manufacturer’s protocol (Fluidigm, 101-0804 B1), labeled with 36 metal-conjugated antibodies in ‘FoxP3 permeabilization buffer’ (eBioscience, 00-8333) with 1% of FBS (Hyclone, Cat# 7207) for 12 hours at 4°C at a concentration of up to 3 million cells per 300 μl of antibody cocktail, followed by twice washing with ‘FoxP3 permeabilization buffer’. Cells were then incubated with the DNA intercalator Iridium (Fluidigm, 201192A) at a dilution of 1:4,000 in 2% paraformaldehyde (Electron Microscopy Sciences, 15714) in DPBS (Corning, 21-031-CV) at RT for 1 hr. Mass cytometry data were acquired on a Fluidigm Hyperion instrument. Flow CyTOF data analyses of endocrine and immune cell composition were performed using the Cytobank implement (https://www.cytobank.org/).

### Imaging Mass Cytometry

Imaging mass cytometry was performed as described (Wang et al., 2019). Tissue slides were labeled with 31 metal-conjugated antibodies, then ablated by a UV-laser, and the resulting plumes of particles were carried to a mass spectrometer for signal detection. Cell segmentation was performed with Vis image analysis software (Visiopharm). Noise was first removed from all images with a 3×3 pixel median filter, followed by nuclear object detection using a polynomial local linear parameter-based blob filter applied to the Iridium-193 DNA channel of each region of interest (ROI). Nuclear objects less than 10μm^2^ were filtered out, then the remaining objects expanded to a maximum of 7 pixels before exporting the mean channel intensities for further analysis. Each ROI was individually z-score normalized prior to cluster analysis, which was performed with Phenograph using the five hormone channels (C-peptide, Ghrelin, Glucagon, Somatostatin, and PP) as input, and a nearest neighbor setting (k) of 200. Endocrine cell types were assigned according to expression of their canonical hormones, and quantified.

### Immunofluorescence Staining for P-CREB

Paraffin embedded fixed pancreatic sections affixed to glass slides were dewaxed, and heat-induced epitope retrieval was performed using citrate buffer. Sections were blocked with 5% donkey serum in the presence of 0.1% Triton followed by overnight incubation with primary antisera at 4 degrees (pCREB, Cell Signaling; Glucagon, Abcam). After extensive washing, sections were incubated with secondary antisera (Jackson ImmunoResearch) for 1 hour at room temperature. Confocal imaging was performed on a Leica TCS SP8 inverted microscope. Using ImageJ to apply a constant threshold across all samples, 500 endocrine cells per donor were manually counted and pCREB and glucagon expression was evaluated. The data were expressed as percent of glucagon and pCREB dual positive cells compared to total glucagon positive cells in control and GADA+ donors. Unpaired t-test was used to compare the groups.

## QUANTIFICATION AND STATISTICAL ANALYSIS

Insulin and glucagon data are presented as the mean ± standard error (SE). Area under curve (AUC) for each intervention was calculated by case, and statistical analysis was then performed by unpaired t-test with Welch’s correction. In appropriate cases, significant differences between groups were determined by ANOVA with post hoc analysis using Dunnett’s multiple comparison test (Dunnett, 1980). *p* ≤ 0.05 was considered significant. IGOR data analysis software (Kline, 2006) (Wavemetrics, Lake Oswego, OR) was used to measure the total insulin or glucagon secretion by calculating the integral under the insulin or glucagon curves, respectively (presented in relative units).

## KEY RESOURCES TABLE

This Table is under construction using

https://star-methods.com/

The KRT reference ID is KRT616708abe4f3a

