## Supplemental Figures S1-S4 and Tables S1 and S2 for "Alpha cell dysfunction in early type 1 diabetes"

### Figure S1

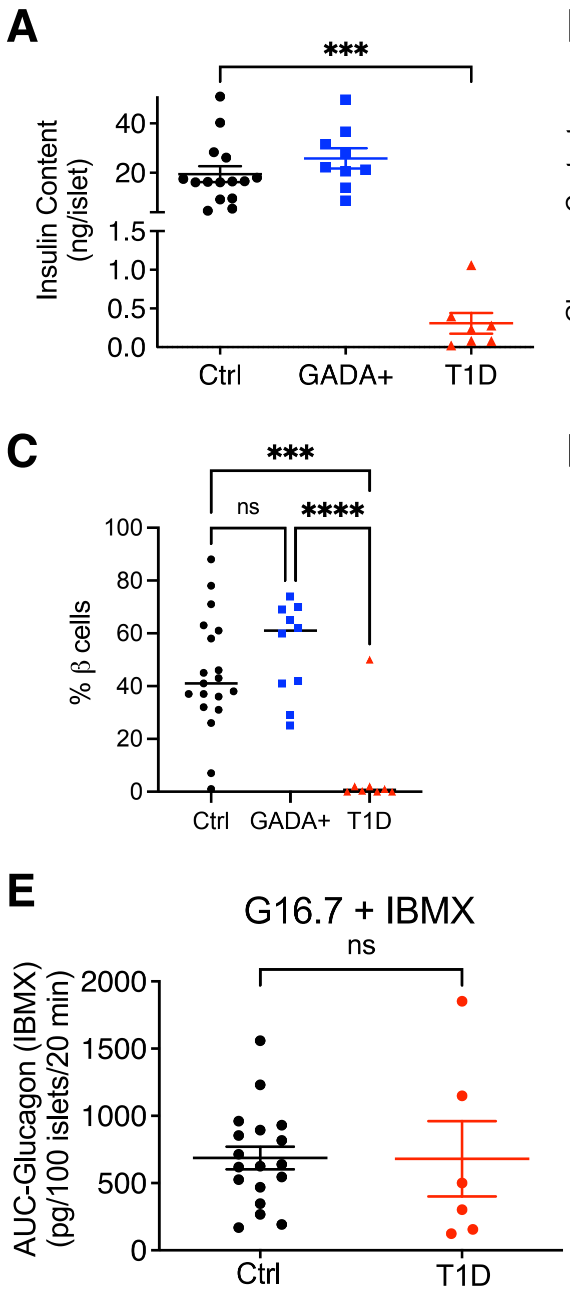
A
***
Insulin Content
(ng/islet)
40
20
1.5
1.0
0.5
0.0
Ctrl
GADA+
T1D
B
ns
ns
Glucagon Content
(pg/islet)
3000
2000
1000
0
Ctrl
GADA+
T1D
C
***
ns
****
% β cells
100
80
60
40
20
0
Ctrl
GADA+
T1D
D
ns
ns
ns
% α cells
100
80
60
40
20
0
Ctrl
GADA+
T1D
E
G16.7 + IBMX
ns
AUC-Glucagon (IBMX)
(pg/100 islets/20 min)
2000
1500
1000
500
0
Ctrl
T1D

### Figure S2

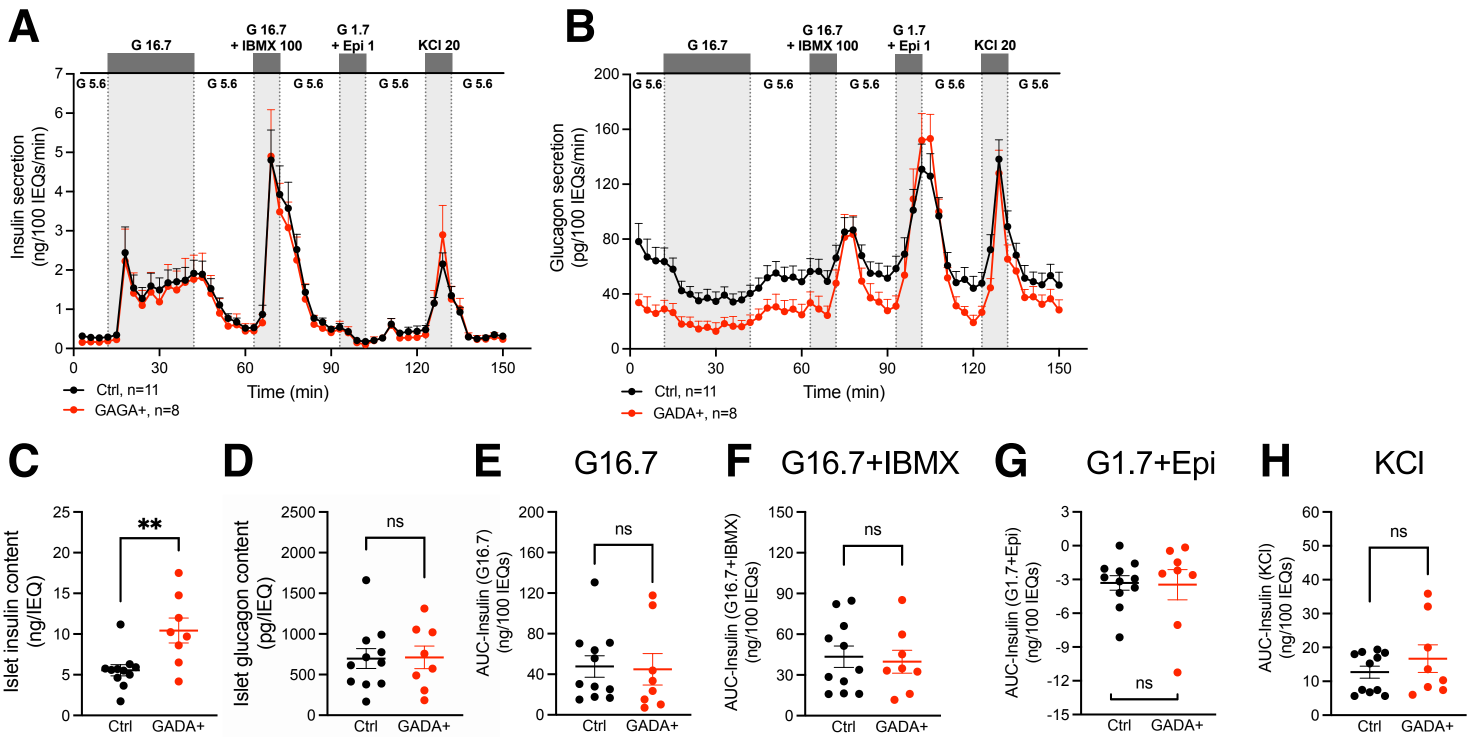
A
G 16.7
G 16.7 + IBMX 100
G 1.7 + Epi 1
KCl 20
G 5.6
Insulin secretion (ng/100 IEQs/min)
Time (min)
Ctrl, n=11
GAGA+, n=8
B
Glucagon secretion (pg/100 IEQs/min)
Ctrl, n=11
GADA+, n=8
C
Islet insulin content (ng/IEQ)
**
D
Islet glucagon content (pg/IEQ)
ns
E
G16.7
AUC-Insulin (G16.7) (ng/100 IEQs)
F
G16.7+IBMX
AUC-Insulin (G16.7+IBMX) (ng/100 IEQs)
G
G1.7+Epi
AUC-Insulin (G1.7+Epi) (ng/100 IEQs)
H
KCl
AUC-Insulin (KCl) (ng/100 IEQs)
Ctrl
GADA+

### Figure S3

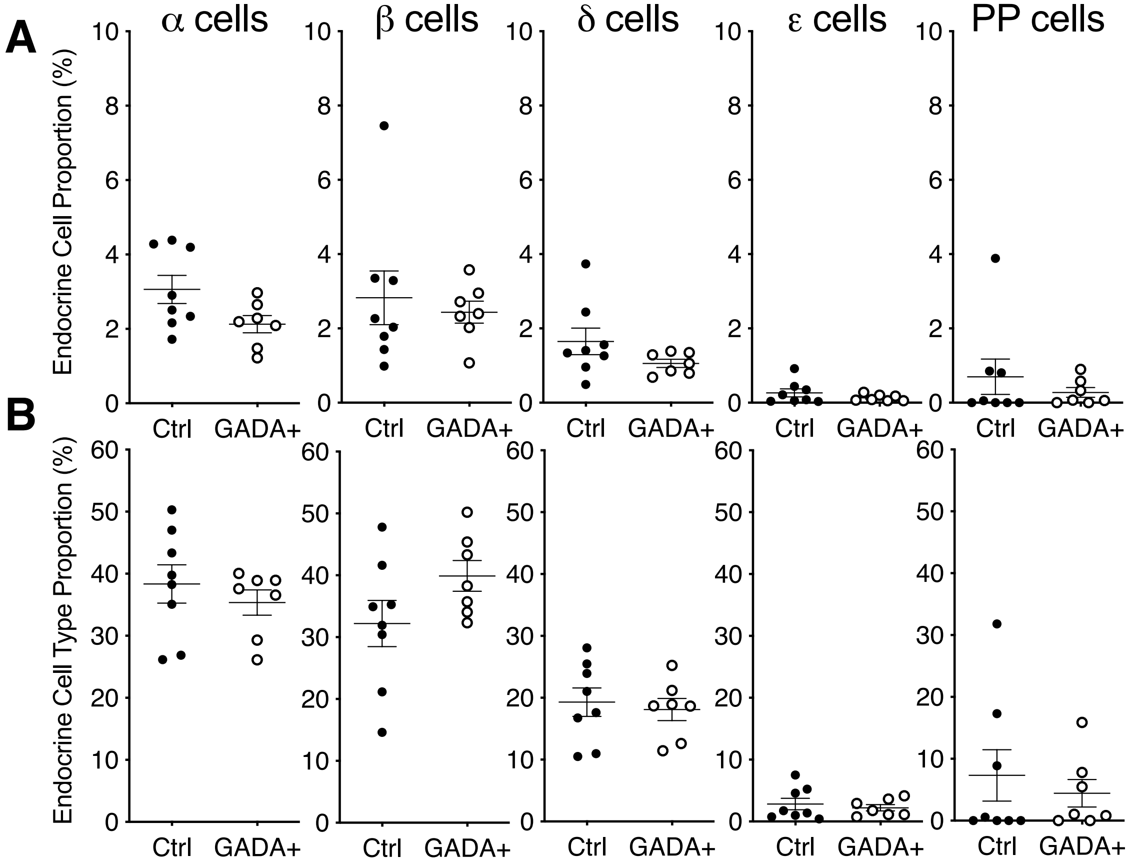
A
B
α cells
β cells
δ cells
ε cells
PP cells
Endocrine Cell Proportion (%)
Endocrine Cell Type Proportion (%)
Ctrl
GADA+
0
2
4
6
8
10
10
20
30
40
50
60

### Figure S4

#### CD4 T cells

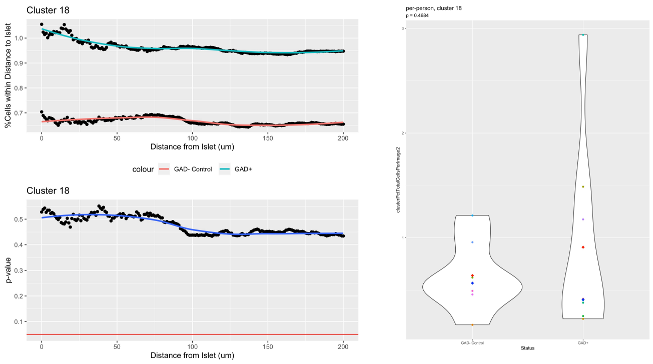


#### Ki67-positive macrophages

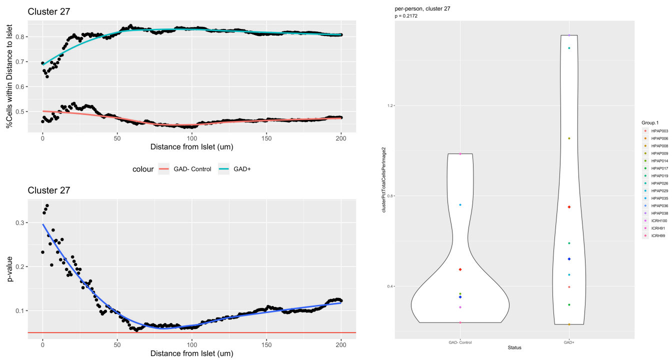


#### CD8 T cells

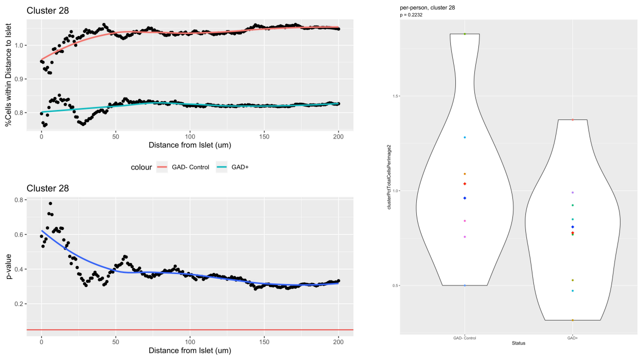


#### Ki67-negative macrophages

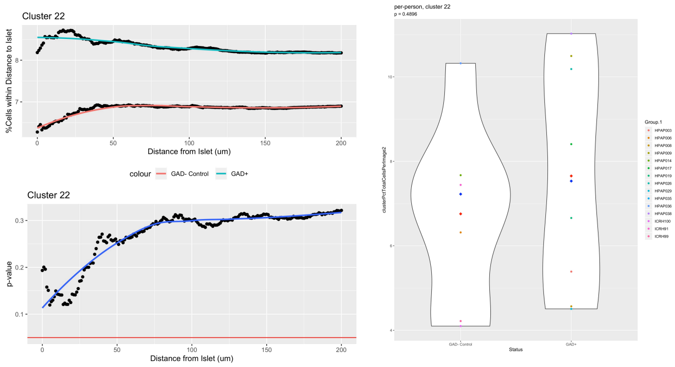


#### B cells

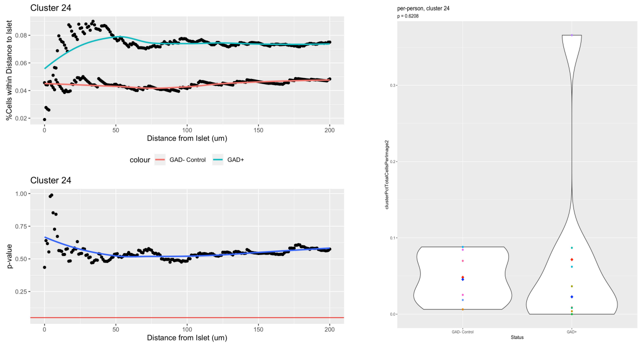

### Table S1

| ID | Recovery Center | Sex | Age (Years) | Race | BMI | Medical History | T1D Duration | AutoAb | HbA1c | C-peptide (ng/mL) | Perifusion (Penn) | Perifusion (Vanderbilt) | Flow CyTOF | IMC | scRNAseq | pCREB Analysis |
|---|---|---|---|---|---|---|---|---|---|---|---|---|---|---|---|---|
| **Control Donors** | | | | | | | | | | | | | | | | |
| HPAP-012 | nPod | F | 18 | Caucasian | 29.6 | — | — | — | 4.5 | 4.1 | x | x | x | x | | |
| HPAP-018 | Penn | M | 31 | Hispanic | 24.5 | — | — | — | 5.4 | 6.40 | x | x | x | x | | |
| HPAP-022 | Penn | F | 39 | Caucasian | 34.7 | — | — | — | 4.7 | 9.35 | | | x | | x | |
| HPAP-026 | nPod | M | 24 | Caucasian | 20.6 | — | — | — | 4.9 | 0.25 | x | x | x | | x | x |
| HPAP-027 | Penn | F | 31 | Caucasian | 32.7 | — | — | — | 4.4 | 7.06 | x | | x | x | | |
| HPAP-034 | Penn | M | 13 | Caucasian | 18.7 | — | — | — | 5.2 | 12.7 | | | x | | x | |
| HPAP-035 | Penn | M | 35 | Caucasian | 26.9 | — | — | — | 5.2 | 15.90 | x | x | x | x | x | x |
| HPAP-036 | nPod | F | 23 | Caucasian | 16.0 | — | — | — | 5.2 | 1.12 | x | | x | x | x | x |
| HPAP-037 | Penn | F | 35 | Caucasian | 21.9 | — | — | — | 5.3 | 4.75 | x | | x | | x | |
| HPAP-039 | nPod | F | 5 | Caucasian | 16.3 | — | — | — | 6.8 | 1.88 | | | x | | x | |
| HPAP-040 | Penn | M | 35 | Caucasian | 24.0 | — | — | — | 5.4 | 7.01 | x | x | x | | x | |
| HPAP-046 | Penn | M | 19 | African-American | 21.0 | — | — | — | 5.7 | 20.74 | x | x | | | | |
| HPAP-047 | Penn | M | 8 | Caucasian | 16.8 | — | — | — | ND | 1.24 | | | x | | x | |
| HPAP-052 | Penn | M | 27 | African-American | 38.7 | — | — | — | 5.2 | 4.07 | x | x | x | | | |
| HPAP-054 | Penn | F | 40 | Caucasian | 30.0 | — | — | — | 4.8 | 6.38 | x | x | x | | | |
| HPAP-056 | Penn | M | 33 | Caucasian | 32.9 | — | — | — | 5.6 | 14.41 | x | x | x | | | |
| HPAP-059 | Penn | M | 35 | Caucasian | 38.0 | — | — | — | 5.1 | 8.18 | x | x | x | | | |
| HPAP-074 | Penn | F | 40 | Caucasian | 36.9 | — | — | — | 6.3 | 4.25 | x | | x | | | |
| HPAP-075 | Penn | M | 35 | Caucasian | 27.5 | — | — | — | 6.0 | 11.97 | x | | x | | | |
| HPAP-080 | nPod | M | 22 | African-American | 35.7 | — | — | — | 5.4 | 15.35 | x | x | x | | | |
| ICRH91* | Penn | F | 35 | Caucasian | 23.6 | — | — | — | 4.6 | N/A | x | | | x | | |
| ICRH99* | Penn | M | 17 | Caucasian | 25.6 | — | — | — | 5.0 | N/A | x | | | x | | |
| ICRH100* | Penn | M | 30 | Caucasian | 22.4 | — | — | — | 5.3 | N/A | x | | | x | | |
| **GADA+ Donors** | | | | | | | | | | | | | | | | |
| HPAP-003 | nPod | M | 29 | Caucasian | 24.5 | — | — | GADA+ | 5.6 | 9.00 | x | x | x | x | | x |
| HPAP-008 | nPod | F | 24 | Caucasian | 31.9 | — | — | GADA+ | 5.2 | 27.05 | x | x | x | x | | x |
| HPAP-017 | nPod | M | 30 | Caucasian | 23.7 | — | — | GADA+ | 5.5 | 3.71 | x | x | x | x | | x |
| HPAP-019 | nPod | M | 22 | Caucasian | 29.8 | — | — | GADA+ | 5.2 | 8.82 | x | x | x | x | | x |
| HPAP-024 | nPod | M | 18 | Caucasian | 24.3 | — | — | GADA+ | 5.5 | 5.6 | | | x | | x | |
| HPAP-029 | nPod | M | 23 | Caucasian | 28.6 | — | — | GADA+ | 5.3 | 3.83 | x | x | x | x | x | |
| HPAP-038 | nPod | M | 13 | Caucasian | 18.3 | — | — | GADA+ | 5.7 | 8.29 | x | x | x | x | x | x |
| HPAP-045 | nPod | F | 27 | Caucasian | 26.2 | — | — | GADA+ | 5.2 | 1.70 | x | x | x | | x | |
| HPAP-049 | nPod | M | 29 | African-American | 32.7 | — | — | GADA+ | 5.4 | 6.15 | x | x | x | | x | |
| HPAP-050 | nPod | F | 22 | Hispanic | 29.0 | — | — | GADA+ | 5.1 | 3.79 | x | | x | | x | |
| **T1D Donors** | | | | | | | | | | | | | | | | |
| HPAP-002 | nPod | M | 26 | Hispanic | 16.4 | T1D | 5 years | — | 9.8 | 0.51 | x | | x | | | |
| HPAP-015 | nPod | M | 29 | Caucasian | 22.0 | T1D | 7 years | mIAA+ | ND | 0.03 | x | | x | | | |
| HPAP-020 | nPod | M | 14 | Caucasian | 13.3 | T1D | 0 (Recent) | GADA+, IA-2+, mIAA+, ZnT8+ | ND | 0.37 | x | | x | | | |
| HPAP-021 | nPod | F | 13 | Caucasian | 21.4 | T1D | 7 years | mIAA+ | ND | <0.02 | x | | x | | | |
| HPAP-055 | Penn | M | 24 | Hispanic | 27.9 | T1D | 7 years | GADA+, IA-2+, mIAA+, ZnT8+ | 10.8 | <0.02 | x | | x | | | |
| HPAP-071 | nPod | F | 12 | Caucasian | 15.4 | T1D | 3 years | IA-2+ | 9.8 | 0.06 | x | | | | | |

### Table S2

| Gene Symbol | Description | $log_2FC$ | padj |
|---|---|---|---|
| MRLN | myoregulin | -2.575 | 0.002 |
| PSMB10 | proteasome 20S subunit beta 10 | -2.591 | 0.002 |
| SAMD11 | sterile alpha motif domain containing 11 | -0.159 | 0.004 |
| G6PC2 | glucose-6-phosphatase catalytic subunit 2 | -2.775 | 0.005 |
| SCD5 | stearoyl-CoA desaturase 5 | -0.205 | 0.005 |
| GPM6A | glycoprotein M6A | -0.081 | 0.005 |
| TCIM | transcriptional and immune response regulator | -0.084 | 0.005 |
| NPM3 | nucleophosmin/nucleoplasmin 3 | -1.702 | 0.005 |
| PDX1 | pancreatic and duodenal homeobox 1 | -0.118 | 0.005 |
| MRPL52 | mitochondrial ribosomal protein L52 | -1.497 | 0.005 |
| SIX3-AS1 | SIX3 antisense RNA 1 | -0.078 | 0.008 |
| PKIB | cAMP-dependent protein kinase inhibitor beta | -2.296 | 0.008 |
| GADD45GIP1 | GADD45G interacting protein 1 | -0.752 | 0.010 |
| NEIL2 | nei like DNA glycosylase 2 | -1.001 | 0.010 |
| FFAR4 | free fatty acid receptor 4 | -0.094 | 0.010 |
| C2orf76 | chromosome 2 open reading frame 76 | -2.329 | 0.011 |
| DLK1 | delta like non-canonical Notch ligand 1 | -0.069 | 0.011 |
| TMEM99 | KRT10 antisense RNA 1 | -0.222 | 0.011 |
| SDHAF3 | succinate dehydrogenase complex assembly factor 3 | -1.848 | 0.011 |
| PCDH7 | protocadherin 7 | -0.079 | 0.015 |
| SEMA6A | semaphorin 6A | -0.153 | 0.020 |
| SNCA | synuclein alpha | -1.379 | 0.021 |
| MAFA | MAF bZIP transcription factor A | -0.149 | 0.021 |
| TGFBR3 | transforming growth factor beta receptor 3 | -0.094 | 0.024 |
| HIBADH | 3-hydroxyisobutyrate dehydrogenase | -1.754 | 0.024 |
| C11orf74 | intraflagellar transport associated protein | -1.929 | 0.024 |
| MRPL24 | mitochondrial ribosomal protein L24 | -1.516 | 0.024 |
| SP110 | SP110 nuclear body protein | -0.094 | 0.024 |
| RTL8C | retrotransposon Gag like 8C | -1.547 | 0.024 |
| PPP1R11 | protein phosphatase 1 regulatory inhibitor subunit 11 | -1.506 | 0.024 |
| STX8 | syntaxin 8 | -1.069 | 0.024 |
| ISOC1 | isochorismatase domain containing 1 | -1.570 | 0.027 |
| TUBB2B | tubulin beta 2B class IIb | -0.165 | 0.027 |
| MOSPD3 | motile sperm domain containing 3 | -0.909 | 0.028 |
| CALD1 | caldesmon 1 | -0.118 | 0.028 |
| TMEM126A | transmembrane protein 126A | -0.985 | 0.028 |
| HSPBP1 | HSPA (Hsp70) binding protein 1 | -1.345 | 0.030 |
| FAM229B | family with sequence similarity 229 member B | -1.054 | 0.030 |
| TMEM126B | transmembrane protein 126B | -1.692 | 0.030 |
| ZNF609 | zinc finger protein 609 | 1.287 | 0.037 |
| S100A11 | S100 calcium binding protein A11 | -1.412 | 0.040 |
| AMN1 | antagonist of mitotic exit network 1 homolog | -1.111 | 0.040 |
| LINC02182 | long intergenic non-protein coding RNA 2182 | -0.177 | 0.040 |
| PYCR2 | pyrroline-5-carboxylate reductase 2 | -1.587 | 0.040 |
| RAB3C | RAB3C, member RAS oncogene family | -1.338 | 0.040 |
| HMGN4 | high mobility group nucleosomal binding domain 4 | -1.846 | 0.043 |
| CCND1 | cyclin D1 | -0.135 | 0.043 |
| BORCS8 | BLOC-1 related complex subunit 8 | -1.480 | 0.043 |
| CMSS1 | cms1 ribosomal small subunit homolog | -1.601 | 0.044 |
| TSHZ2 | teashirt zinc finger homeobox 2 | -0.091 | 0.044 |
| SIX3 | SIX homeobox 3 | -0.099 | 0.044 |
| MGST3 | microsomal glutathione S-transferase 3 | -0.687 | 0.045 |
